# Time-efficient relaxation measurements by ^31^P MR fingerprinting in human brain at 7T

**DOI:** 10.1101/2022.05.23.493067

**Authors:** Song-I Lim, Mark Stephan Widmaier, Daniel Wenz, Jiang Yun, Lijing Xin

## Abstract

**Purpose:** The goal of the study is to develop ^31^P spectroscopic MRF at 7T to measure *T*_1_ and *T*_2_ relaxation times simultaneously and to compare time efficiency and test-retest reproducibility of MRF with conventional inversion recovery and multi-TE methods.

**Methods:** A ^31^P MRF scheme was designed based on a balanced steady-state free precession type sequence. Dictionary was generated using the Bloch equations. *B*_0_ map was acquired experimentally and incorporated into the dictionary. 7 phantoms with different *T*_1_ and *T*_2_ relaxation times were prepared for MRF validation. Simulations were performed to evaluate estimation bias. 7 volunteers were scanned twice using both MRF and the conventional methods to evaluate the reproducibility.

**Results:** In phantom measurements, *T*_1_ and *T*_2_ values between MRF and conventional methods demonstrated a good agreement with Pearson’s correlation coefficients of 0.99 and 0.97, respectively. In *in vivo* experiments, estimated *T*_1_ by MRF were in good agreement with those measured by the inversion recovery and in the literature. On the other hand, estimated *T*_2_ values by MRF were shorter than those measured by the multi-TE method. ^31^P MRF method can reduce the acquisition time by 15 min providing less than 10% of mean CV for *T*_1_ estimations and less than 20% of mean CV for *T*_2_ estimations of metabolites.

**Conclusion:** Our results shows the feasibility of simultaneous *T*_1_ and *T*_2_ measurement of ^31^P metabolites in human brain using MRF at 7T. High reproducibility can be achieved especially for *T*_1_ measurement with 40% time reduction over conventional methods.

## Introduction

^31^P magnetic resonance spectroscopy (MRS) has been widely used to measure energy metabolism in the human brain and muscle (1; 2). The knowledge of *T*_1_ and *T*_2_ relaxation times of metabolites is essential to optimize the measurement parameters of ^31^P MRS acquisition and quantification of metabolite concentration. Moreover, alterations in *T*_1_ and *T*_2_ relaxation times could be linked to changes in cellular and molecular environments caused by pathophysiolgical conditions (3). However, due to the low signal-to-noise (SNR) ratio, measurement of *T*_1_ and *T*_2_ relaxation times suffers from long acquisition duration using conventional measurement methods. Furthermore, it is challenging to measure the *T*_2_ relaxation times of scalar coupled metabolites such as adenosine triphosphate (ATP) with varying echo time (TE) due to the complex J-coupling interaction *in vivo* (4). So far, only the limited number of studies have reported *T*_1_ and *T*_2_ relaxation times in human brain at 7T (5; 6; 7) and *T*_2_ relaxation time of *β*ATP has not been reported yet.

Magnetic resonance fingerprinting (MRF) is a novel technique for fast and quantitative multiparametric acquisition. By changing sequence parameters such as TR, TE, and/or flip angles (FAs), MRF signal evolution can provide a unique signal pattern sensitive to tissue parameters (such as *T*_1_ and *T*_2_ relaxation times, and proton density), *B*_0_, and *B*_1_ fields (8). To extract these parameters, the measured signal pattern is then matched with the predefined dictionary generated by the Bloch simulations. MRF has been extensively studied to further reduce scanning duration and computational load for dictionary matching and to improve accuracy (9; 10; 11; 12). Even though MRF technique has proven its robustness and time-efficiency in human MRI studies, there are sparse studies focusing on spectroscopic MRF for both proton and X-nuclei (13; 14; 15).

^31^P creatine kinase MRF (CK-MRF) was demonstrated in resting rat skeletal muscle to measure the metabolic rate of CK, *T*_1_ relaxation time of PCr and PCr/*γ*ATP ratio and to shorten scanning duration (15). In this study, *T*_2_ of PCr, *T*_1_ and *T*_2_ of *γ*ATP and the field inhomogeneities were fixed to reduce the complexity of parameter estimation (15).

In the current study, we aim to investigate the feasibility of ^31^P spectroscopic MRF to measure relaxation times of ^31^P metabolites in the human brain at 7T. The performance of the ^31^P MRF measurement was first evaluated by simulations, then validated by *in vitro* experiments, and finally its time efficiency and reproducibility were compared with the conventional inversion recovery and multi-TE methods *in vivo*.

## Methods

### MRF sequence design

The ^31^P-MRF pulse sequence design is based on an 1D balanced steady-state free precession (bSSFP) type of sequence as the initial framework (8). The MRF pulse train contains 0 - 180^*°*^phase alternating RF pulses with variable FAs following a sinusoidal pattern and a fixed TR (18.9 ms). 1D localizazion was achieved using an 1.5 ms Shinnar-le-Roux (SLR) slice selective excitation pulse with a bandwidth (BW) of 6.6 kHz. Each excitation is followed by a 16.66 ms of acquisition time with 256 data points.

Figure 1 shows the schematic of the pulse train, which consists of two parts: A) an adiabatic inversion pulse (HS4; 10.24 ms; inversion BW = 4.9 kHz; *γB*_1_ = 0.5 kHz) with 10 ms of inversion delay was followed by 249 FAs. The second inversion pulse with same profile was applied after 200 ms of delay and followed by 270 FAs. Total 519 FAs were transmitted and spoiling gradients were applied after both inversion pulses to remove remaining transverse magnetisation. B) 180 drastic changing excitation FAs enveloped in a sinusoidal shape are followed to increase sensitivity for sensing *B*_1_ and *B*_0_ inhomogeneities (16). Overall, one cycle of MRF acquisition is composed of two inversion pulses with inversion delays, 699 FAs, and 5 s of delay between cycles (21.5 s per cycle).

**Figure 1:**
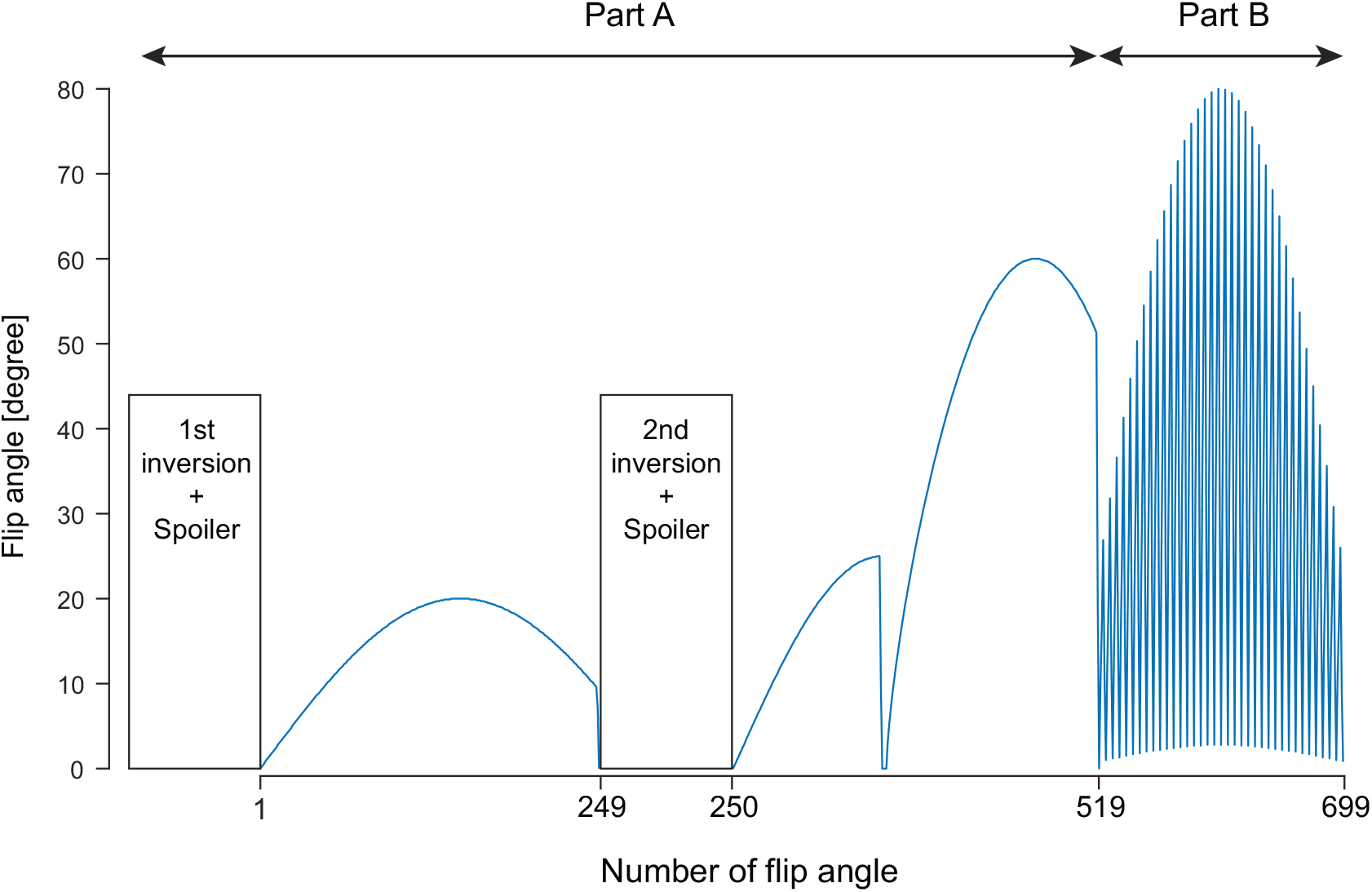
Sequence diagram of ^31^P MRF scheme. One cycle of MRF sequence consists of two parts: part A) with sinusoidal flip angle patterns (519 flip angles) and part B) with drastically changing flip angle patterns sensitive to *B*_0_ and *B*_1_ inhomogeneity. There is a 5 s of delay between cycles.

### Dictionary Simulations

Dictionary was generated based on the Bloch equations. The bSSFP signal profile was simulated and measured experimentally (Figure S1). The effective excitation FA, *ϕ*_*eff*_ (*n*) was found by

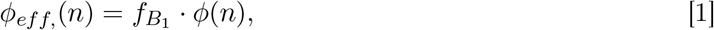

where *ϕ*(*n*) is the *n*th FA in the MRF sequence pattern and the *B*_1_ scaling factor *f*_*B*1_ is

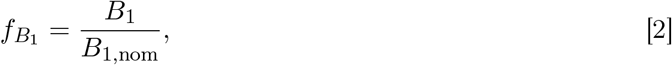

with *B*_1,nom_ the nominal transmit field strength to reach the right FA and *B*_1_ is the actual transmit magnetic field strength seen. Signal evolutions for isochromats ranging from -60 to 60 Hz (1 Hz step) around the respective chemical shift frequency were simulated for each metabolite. The chemical shift of *γ*−, *α*−, *β*ATP and Pi were set to −2.52 ppm, −7.42 ppm, −15.91 ppm and 4.71 ppm relative to that of PCr (0 ppm). A *B*_0_ distribution (−60 Hz to 60 Hz; 1 Hz resolution) was incorporated using the histogram of the measured *B*_0_ map (for *in vitro* and *in vivo* data) with zero mean. Off-resonance effect was included by shifting the mean off-resonance frequency of the *B*_0_ distribution by *f*_*off*_. For ATP resonances, linewidth broadening due to J-coupling was also taken into account using the convolution of the *B*_0_ distribution with Dirac functions separated by the J-coupling constant of each metabolite (J_*βα*_ = 16.1 Hz and J_*βγ*_ = 16.3 Hz) (17). The subject specific signal evolution for one parameter set, was found by the weighted sum of the signal evolutions of each isochromat, given by *B*_0_ distribution, *f*_*off*_, J-coupling effects, and spectral leakage due to discrete Fourier transformation. 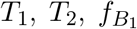 and *f*_*off*_ were free parameters of the dictionary simulation for each metabolite.

### Pattern matching

A diagram of pattern matching steps is illustrated in Figure 2. First, general dictionaries were created for each metabolite with isochromats, *f*_*off*_ (−60 Hz to 60 Hz in 121 steps), and 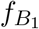 (0.6 to 1.2 in 31 steps). The relaxation parameters *T*_1_ and *T*_2_ for each metabolite were ranging in 16 and 31 steps respectively, according to the values in a supporting table (Table ST1) in a relative or an absolute manner. A subject-specific dictionary was then created for each metabolite out of their general dictionaries as described above. The phase of the resulting complex signal evolutions was corrected by minimizing the imaginary components of part A. All fits were achieved by inner product of the L2 norm of the measured pattern and the dictionary entries. 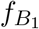 was assumed to be equal for all metabolites. Firstly, 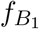 was found by fitting concatenated measured ^31^P MRF signal evolutions of PCr and *γ*ATP using both part A and B with the concatenated dictionaries. Secondly, for each metabolite, the part A and B of the measured ^31^P MRF signal evolutions was used to fit *f*_*off*_. In the last step, part A was used to fit the relaxation parameters of each metabolite. 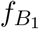 and *f*_*off*_ were set to the prior estimates. All non-assigned parameters in each fitting step were considered as free parameters, even if their estimates were not used in a subsequent fitting step.

**Figure 2:**
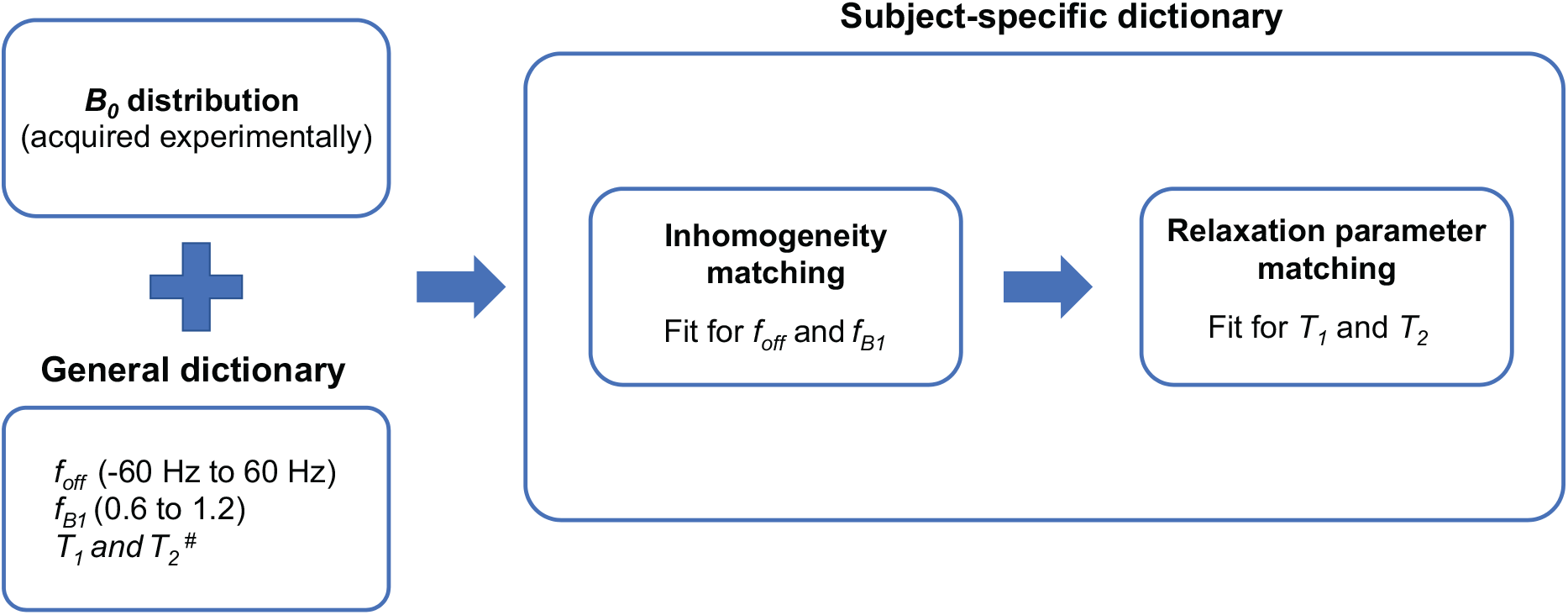
Pattern matching steps. After generating general dictionaries using different 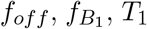 and *T*_2_ values, *B*_0_ distributions are incorporated in order to create subject-specific dictionaries. *f*_*off*_ and 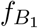 matching is preceded by relaxation parameter matching step. *T*_1_ and *T*_2_ of each metabolite are fitted independently. ^#^: *T*_1_ and *T*_2_ ranges are presented in a supporting table (Table ST1)

### Simulation validation

To investigate the impact of the three step matching procedure on the accuracy of relaxation time estimation, the matching errors (MEs) of 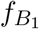, *T*_1_ and *T*_2_ values depending on different ground truth values of 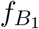 and *f*_*off*_ were evaluated using simulation. The ME was calculated as the relative difference between the estimated parameters and the ground truth values used in the simulation. 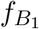 and *f*_*off*_ were ranging from 0.8 to 1.2 and from -10 to 10 Hz, respectively, which were altered independently. 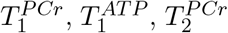 and 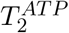 were set to 3.8 s, 1.0 s, 60 ms, and 30 ms respectively. Only a single isochromat was considered as an off-resonance distribution. A Monte-Carlo simulation was performed to further evaluate the matching procedure, noise robustness, and error due to quantification steps. 1000 pairs of PCr and *γ*ATP signal evolutions were created with 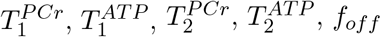 and 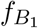 values, which were initialised uniformly and randomly in the range [3.3;4.3] s,[0.8; 1.2] s, [40; 80] ms, [25; 45] ms, [0.8; 1.2], and [-5; 5] Hz in order. *f*_*off*_ and 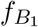 were set the same for both PCr and *γ*ATP signal evolutions. For both simulation evaluations, the same dictionary was used with *f*_*off*_ ranging from −15 Hz to 15 Hz in 31 steps in addition to the chemical shift and 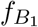 from 0.7 to 1.3 in 31 steps. The relaxation parameters *T*_1_ and *T*_2_ for each metabolite were ranging in 29 and 26 steps, respectively as described in Table ST1.

### *In vitro* validation

7 phantoms with 100 mM Pi and different concentrations of MnCl_2_ and a contrast agent (gadobutrol, Gadovist; Bayer Healthcare Medical Care, Berlin, Germany) were prepared to validate the ^31^P-MRF sequence. All chemicals, otherwise stated, were ordered from Sigma Aldrich. 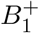 was modulated by changing the transmit voltage. The inversion recovery method was used to measure *T*_1_ values in each phantom (TR = 30 s; inversion delay = 20, 50, 80, 150, 300, 500, 750, 1000, 1500, 2000, 3000, 4000, 5000, 7500, 10000, 12500, 15000, 20000 ms; average = 1; vector size = 2048; slice thickness = 30 mm). The sSPECIAL was used to measure *T*_2_ with varying TE (TR = 30 s; TE = 20, 25, 30, 40, 50, 60, 75, 100, 150, 200, 250, 300, 400 ms; vector size = 2048; VOI = 160×160×30 mm^3^). MRF sequence was performed using the parameters described in Figure 1 (average = 4; 1 dummy scan; slice thickness = 30 mm). *B*_0_ field map acquired by a gradient echo sequence (TR/TE1/TE2 = 9.6/3.37/8.37 ms; flip angle = 5^*°*^; matrix = 256×256; slice thickness = 30 mm). Two general dictionaries were created with 31 steps of *T*_2_ values and 31 steps of *T*_1_ values to cover the range of relaxation parameters. Lower bounds of the dictionaries and step sizes for the relaxation parameter matching are presented in Table ST1. After generating individual dictionaries and incorporating the *B*_0_ maps, fitting procedure (step 1 and 2) were started for 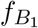 and *f*_*off*_ matching simultaneously.

### *In vivo* experiments

All MR acquisitions were performed on a 7T/68cm MR scanner (Magnetom, Siemens Medical Solutions, Erlangen, Germany) with a two-loop ^1^H and a single loop ^31^P coil. *In vivo* experiments were performed in 7 healthy subjects (3 female and 4 males; age 19-27 years) who provided written informed consent under the approval of the Swiss cantonal ethics committee. Each volunteer was scanned twice to assess test-retest reproducibility of conventional measurement methods and ^31^P MRF. The order of sequence application was randomized to avoid systematic errors. *B*_0_ shimming was performed using the 3D map shim. Transmit voltage was calibrated for setting flip angle correctly. The linewidth of PCr was measured before the application of each sequence. The slice selective SLR pulse (1.5 ms, BW = 6.6 kHz) is used for the ^31^P inversion recovery and MRF sequence. *T*_1_ relaxation time was measured using inversion recovery method based on a slice selective FID sequence (TR = 4 s; inversion delay = 60, 200, 400, 670, 1590, 3000 ms; slice thickness = 20 mm; average = 16; 1 dummy scan; BW = 6000 Hz). Two reference spectra without inversion pulse were acquired using the same sequence (TR = 20 and 4 s; average = 6 and 16; 1 dummy scan; slice thickness = 20 mm; BW = 8000 Hz). The sSPECIAL sequence was used to measure *T*_2_ relaxation times (TR = 7.5 s; TE = 15, 20, 27, 35, 48, 70, 100, 200 ms; VOI = 160×160×20 mm^3^; varying averages from 8 to 16; 2 dummy scan; BW = 8000 Hz). Total scan duration of *T*_1_ and *T*_2_ measurements for 26 min 20 s. The following parameters were used for MRF experiments: TR/TE = 18.9/9.45 ms, number of flip angles = 699, slice thickness = 20 mm, averages = 83, 1 dummy scan, scan duration = 26 min 36 s). For comparison, the number of averages of MRF was set to achieve the same scan duration as the *T*_1_ and *T*_2_ measurements. A *B*_0_ field map was acquired by a gradient echo sequence (TR/TE1/TE2 = 9.6/3.37/8.37 ms, flip angle = 5^*°*^, matrix = 256×256, slice thickness = 30 mm) to incorporate its *B*_0_ distribution into the dictionary in the post-processing step.

### Data processing

All data processing, simulations, and statistical analysis were performed using MATLAB (MathWorks, Natick, MA, USA). 6 Hz of Gaussian apodization was applied to all FIDs before Fourier transformation using FID-A(18). Phase correction was preceded before ^31^P spectra fitting. The intensity of each ^31^P peaks was fitted by Voigt lineshape (a combination of Gaussian and Lorentzian lineshape). *T*_1_ and *T*_2_ fittings were performed using a least-squares function *lsqcurvefit*. Each fitted parameters were ensured to be within the preset upper and lower bounds. Sub-datasets with different number of averages were generated to evaluate the time efficiency of all methods.

### Statistical analysis

The percentage coefficient of variation (CV)

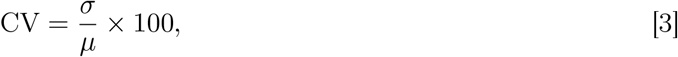

where standard deviation and mean are given as *σ* and *µ*, was used to assess test-retest reproducibility between the two sessions. To compare time efficiency of the two methods, CV changes over the scanning duration was evaluated. The agreement between methods are assessed using Pearson’s correlation (\**p* < 0.05, \*\**p* < 0.01, \*\*\**p* < 0.001).

## Results

### ^31^P MRF scheme validation

MEs induced by varying ground truth 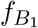 and *f*_*off*_ values are illustrated in Figure 3 (a,d). Fitted *T*_1_ values were not affected neither by 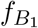 nor *f*_*off*_. One the other hand, fitted *T*_2_ values were underestimated for high 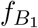 by 4% and in a range between -3 and 3 Hz of *f*_*off*_ by 4%. The SNR dependency of MEs of 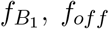, and *T*_1_ and *T*_2_ of PCr and *γ*ATP are presented in Figure 3 (b,c,e,f). Estimated 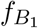 and *f*_*off*_ values were rarely affected by SNR and lower than 5.4% and 1.2%, respectively. For relaxation parameter estimation, *T*_1_ of PCr fitting was more robust to SNR deviation than that of *γ*ATP. *T*_2_ estimation showed similar results for both PCr and *γ*ATP and higher MEs than other parameters.

**Figure 3:**
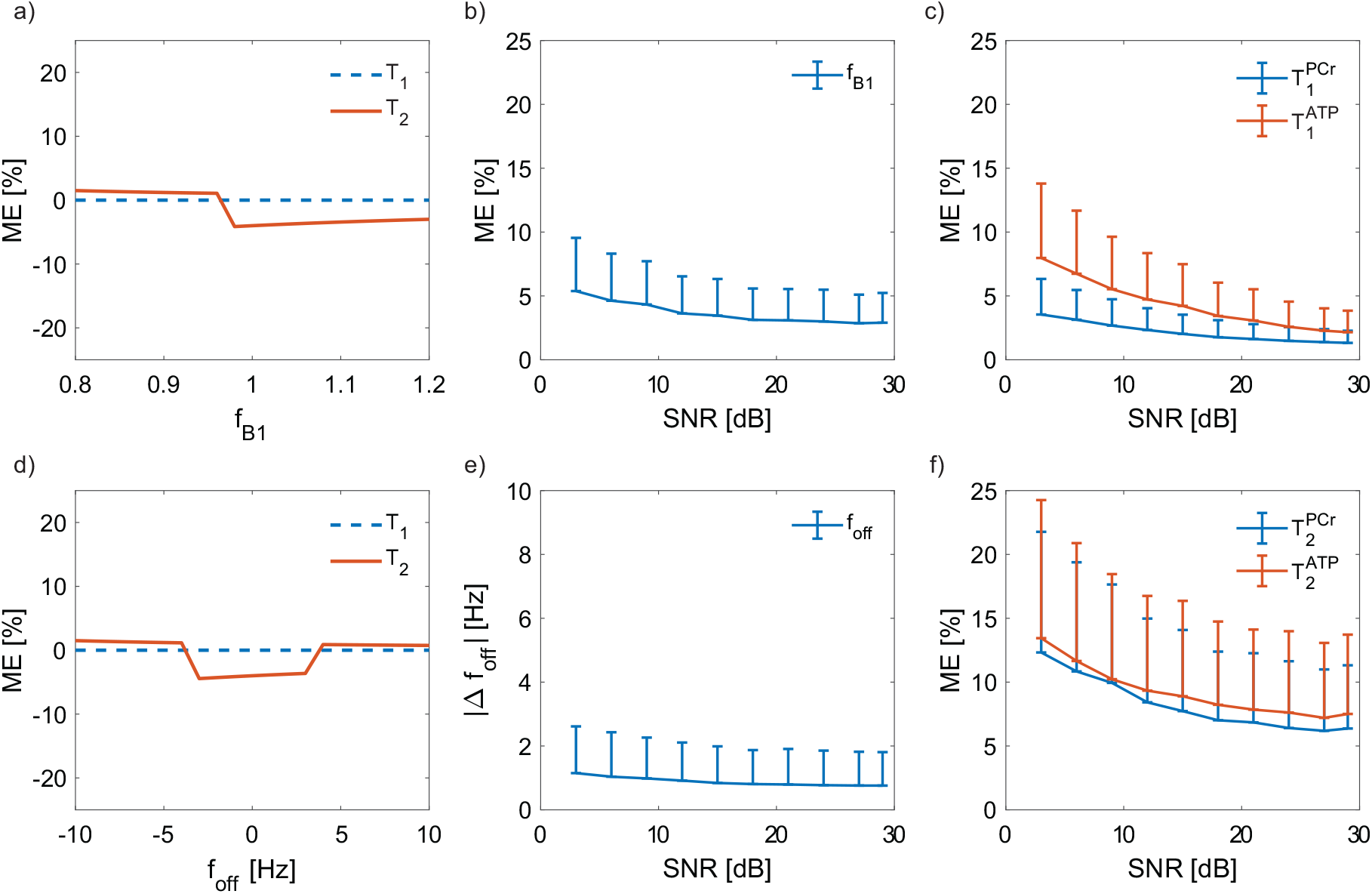
Matching errors (MEs) caused by field inhomogenety and SNR. MEs of *T*_1_ and *T*_2_ of on-resonance peak induced by different 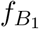 and *f*_*off*_ inputs (a,d). SNR dependency of MEs for 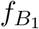, and *T*_1_ and *T*_2_ of PCr and *γ*ATP (c,f). The MEs of *f*_*off*_ is expressed in absolute differences as | Δ*f*_*off*_ | [Hz].

Table 1 shows the comparison of *T*_1_ and *T*_2_ values estimated by conventional methods and MRF. *In vitro* experiments show that *T*_1_ values obtained by the MRF method are in excellent agreement (within 5% of difference) with those by the inversion recovery method. The Pearson’s correlation coefficient for *T*_1_ and *T*_2_ values between MRF and conventional methods were 0.99 and 0.97, respectively as shown in Figure 4. *T*_2_ relaxation times measured by the MRF method are 22% shorter than those acquired by the multi-TE method. To evaluate the effect of transmit voltage on relaxation value estimation, input transmit voltage varies in a range of 55 and 75 V. The matched 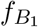 values increased linearly in accordance with the increasing input transmit voltage. Similar to the simulation results, the transmit voltage has minor effects on *T*_1_ and *T*_2_ values. Representative *in vitro* phantom reference image and pattern matching results are presented in Figure S2.

**Table 1:** *In vitro* MRF validation results using 7 phantoms containing different concentrations of gadolinum and MnCl_2_. Measured *T*_1_ and *T*_2_ relaxation times using conventional methods are presented for comparison. To evaluate the effect of 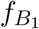, MRF experiments were performed three times using different input voltages (55, 65, and 75 V).

| | Concentrations | | Conventional methods | | U [V] | $^{31}\text{P}$-MRF | | |
| --- | --- | --- | --- | --- | --- | --- | --- | --- |
| | Gado [$\mu\text{M}$] | $\text{MnCl}_2$ [$\mu\text{M}$] | $T_1$ [s] | $T_2$ [ms] | | $T_1$ [s] | $T_2$ [ms] | $f_{B_1}$ |
| P1 | 250 | 2 | 0.86 | 35 | 55 | 0.79 | 30 | 0.91 |
|  |  |  |  |  | 65 | 0.79 | 28 | 0.98 |
|  |  |  |  |  | 75 | 0.79 | 27 | 1.06 |
| P2 | 150 | 2 | 1.18 | 36 | 55 | 1.12 | 36 | 0.89 |
|  |  |  |  |  | 65 | 1.12 | 36 | 0.97 |
|  |  |  |  |  | 75 | 1.12 | 36 | 1.06 |
| P3 | 30 | 0.5 | 3.59 | 77 | 55 | 3.18 | 61 | 0.86 |
|  |  |  |  |  | 65 | 3.18 | 61 | 0.95 |
|  |  |  |  |  | 75 | 3.18 | 61 | 1.01 |
| P4 | 50 | 0.25 | 2.61 | 116 | 55 | 2.67 | 86 | 0.86 |
|  |  |  |  |  | 65 | 2.67 | 86 | 0.94 |
|  |  |  |  |  | 75 | 2.67 | 86 | 1.00 |
| P5 | 0 | 1 | 4.91 | 47 | 55 | 4.51 | 36 | 0.88 |
|  |  |  |  |  | 65 | 4.51 | 36 | 0.97 |
|  |  |  |  |  | 75 | 4.51 | 38 | 1.05 |
| P6 | 15 | 0.01 | 4.51 | 127 | 55 | 4.69 | 115 | 0.94 |
|  |  |  |  |  | 65 | 4.69 | 121 | 1.02 |
|  |  |  |  |  | 75 | 4.69 | 115 | 1.08 |
| P7 | 10 | 0.05 | 5.47 | 109 | 55 | 5.73 | 91 | 0.85 |
|  |  |  |  |  | 65 | 5.73 | 86 | 0.91 |
|  |  |  |  |  | 75 | 5.73 | 86 | 0.98 |

**Figure 4:**
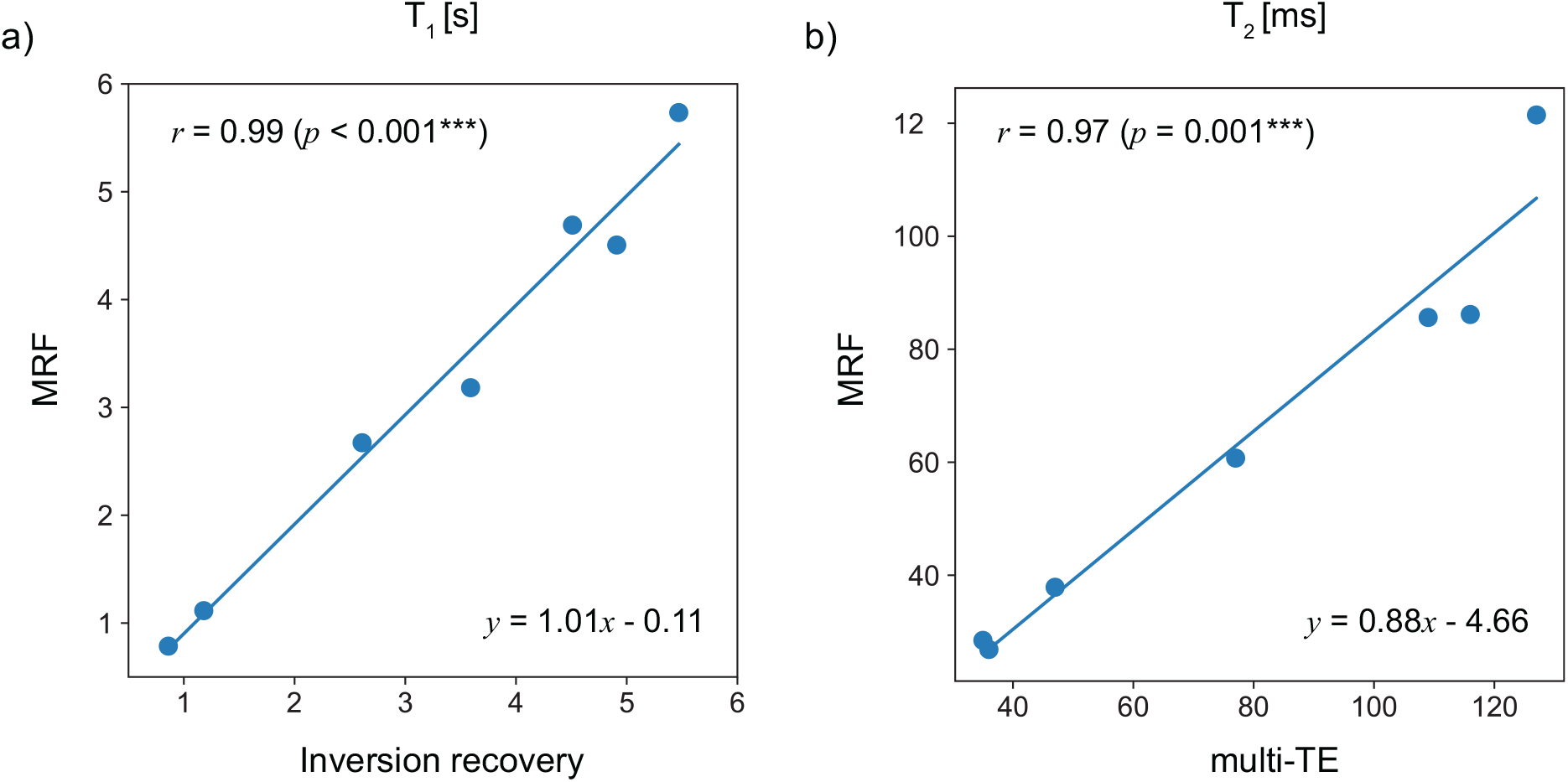
The agreement between MRF and conventional methods. The estimated *T*_1_ and *T*_2_ values of 7 phantoms using MRF are compared with those acquired from inversion recovery (a) and multi-TE (b) methods. The Pearson’s correlation coefficients and *p*-values are presented in the figure (\*\*\**p* < 0.001).

### *In vivo* measurement

The representative *in vivo* MR fingerprints and dictionary matches are presented in Figure 5. The measured fingerprints of all metabolites and corresponding dictionary matches align well each other. *T*_1_ and *T*_2_ fitting curves of conventional methods are shown in Figure 6. Estimated *T*_1_ and *T*_2_ values acquired in our experiments and literature values are provided in Table 2.

**Figure 5:**
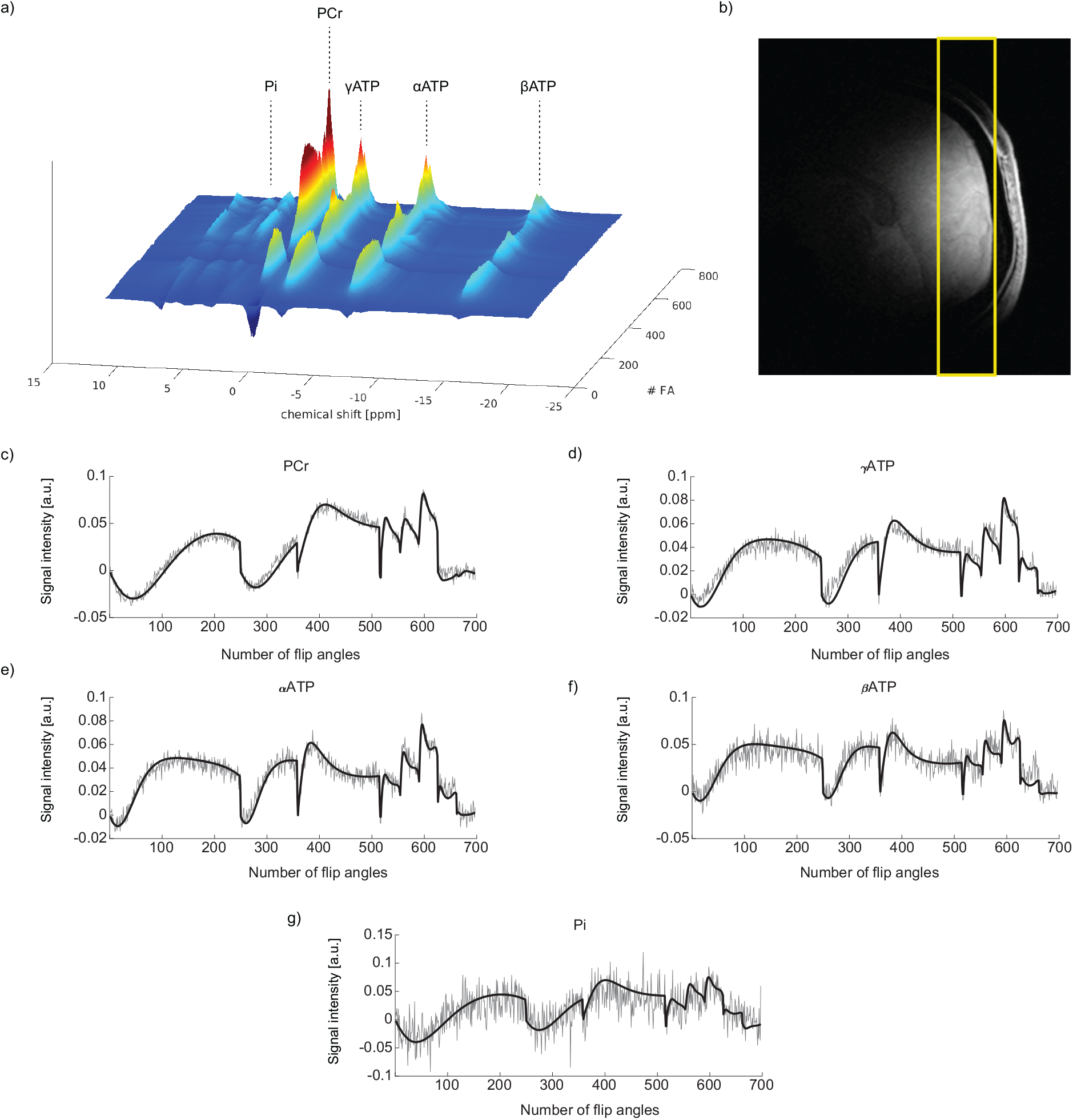
Representative *in vivo* MRF spectra and fingerprint matching results. ^31^P MRF spectra of each FA (83 averages) are illustrated in (a). The representative anatomical image with the slice positioning in yellow is shown in (b). (c-g) present MR fingerprints in gray and their dictionary matches in black for PCr, *γ*ATP, *α*ATP, *β*ATP, and Pi, respectively.

**Figure 6:**
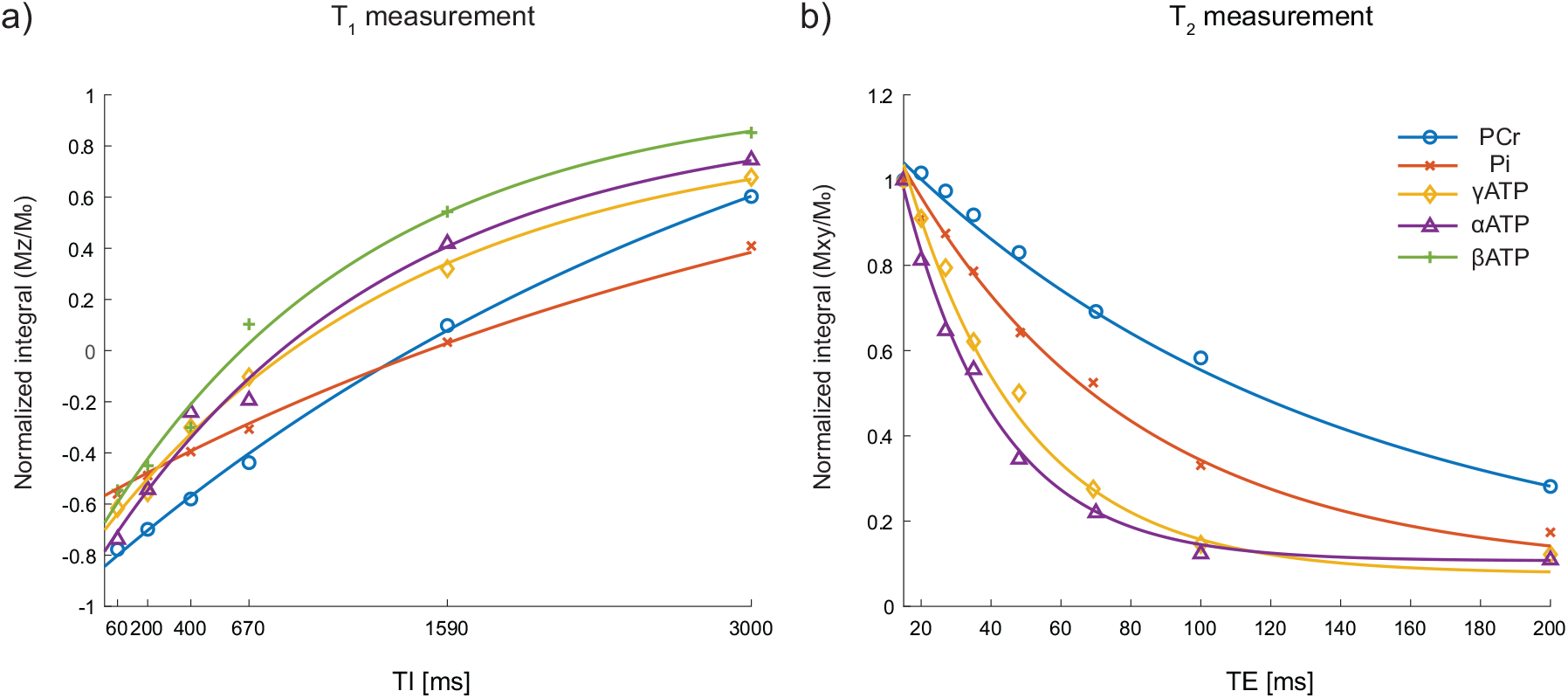
Representative *T*_1_ and *T*_2_ curve fitting results of inversion recovery and multi-TE methods.

**Table 2:** Literature and estimated *T*_1_ and *T*_2_ values of ^31^P metabolites in human brain at 7T using

|  |  | Conventional methods | MRF | References |
| --- | --- | --- | --- | --- |
| $T_1$ [s] | PCr | $3.45 \pm 0.17$ | $3.76 \pm 0.17$ | 3.37 - 3.39 (5; 6) |
| | Pi | $3.88 \pm 0.32$ | $3.89 \pm 0.84$ | 3.19 - 3.70 (5; 6) |
| | $\gamma\text{ATP}$ | $1.33 \pm 0.21$ | $1.11 \pm 0.18$ | 1.27 - 1.70 (5; 6) |
| | $\alpha\text{ATP}$ | $1.26 \pm 0.16$ | $0.88 \pm 0.11$ | 1.26 - 1.35 (5; 6) |
| | $\beta\text{ATP}$ | $1.04 \pm 0.17$ | $0.81 \pm 0.14$ | 1.02 - 1.13 (5; 6) |
| $T_2$ [ms] | PCr | $137.0 \pm 19.4$ | $64.9 \pm 9.0$ | $132.0 \pm 12.8$ (5) |
| | Pi | $64.2 \pm 7.7$ | $46.5 \pm 12.6$ | $86 \pm 2$ (7) |
| | $\gamma\text{ATP}$ | $29.2 \pm 2.3$ | $34.0 \pm 4.5$ | $26.1 \pm 9.6$ (5) |
| | $\alpha\text{ATP}$ | $28.2 \pm 2.9$ | $23.3 \pm 6.3$ | $25.8 \pm 6.6$ (5) |
| | $\beta\text{ATP}$ | - | $25.0 \pm 8.3$ | - |

In Figure 7, measured *T*_1_ and *T*_2_ relaxation times are not biased over different acquisition times are compared between the conventional methods and MRF. *T*_1_ of PCr measured by the inversion recovery method increases with the increasing acquisition time and converges. On the other hand, *T*_2_ values of *γ*- and *α*ATP, and Pi decrease with the increasing number of averages. In general, *T*_1_ and *T*_2_ values estimated by MRF shows over different acquisition durations. Furthermore, standard deviations of MRF determined relaxation times are smaller relative to those of conventional methods especially at short acquisition times, except for *T*_2_ of Pi.

**Figure 7:**
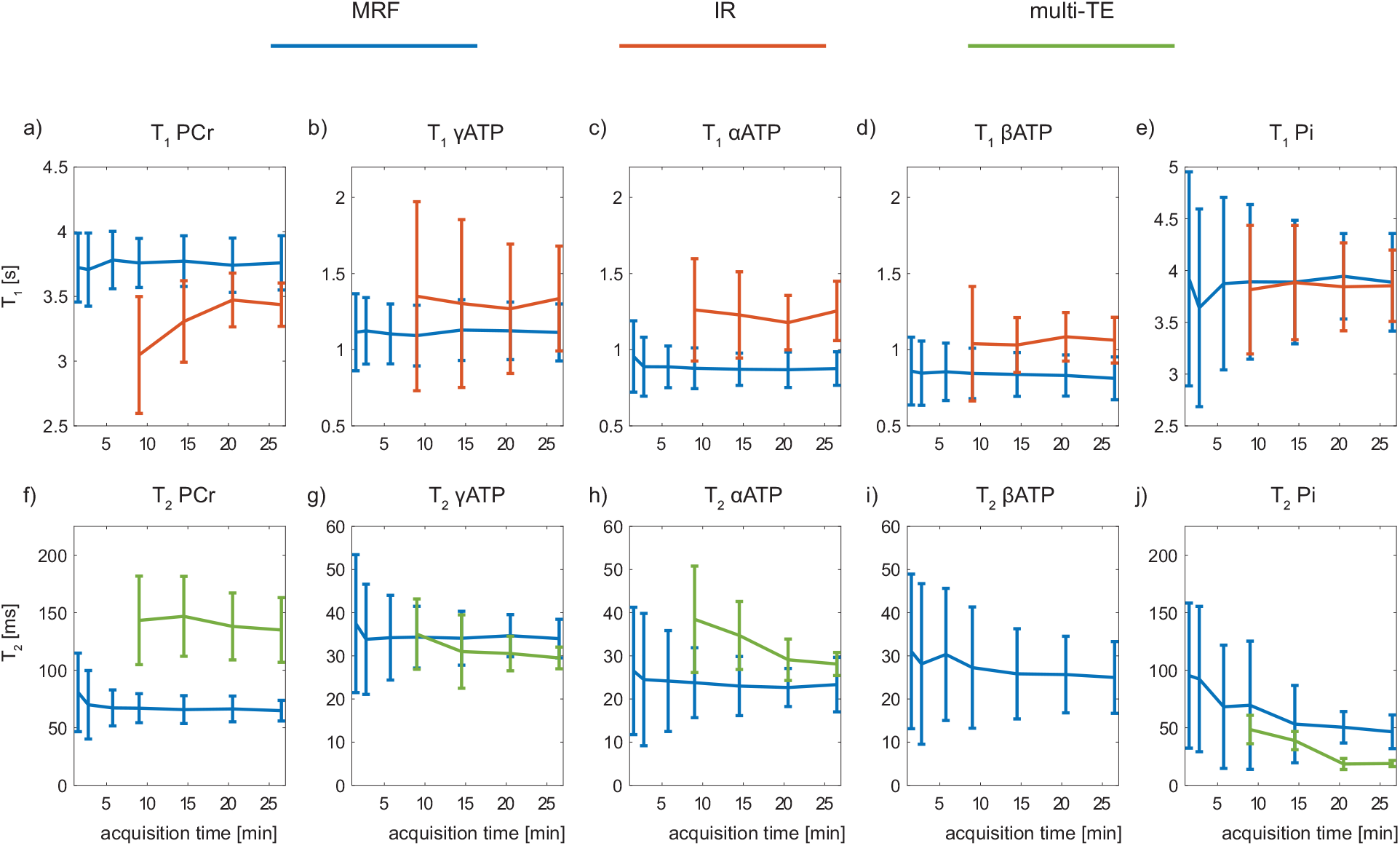
Comparison of estimated mean *T*_1_ and *T*_2_ changes over acquisition time with SDs. Matched *T*_1_ and *T*_2_ by MRF is presented in blue. Fitted *T*_1_ by inversion recovery and *T*_2_ by multi-TE methods are illustrated in orange and green, respectively.

The test-retest reproducibility was evaluated using CV. CVs of estimated *T*_1_ values of the metabolites by MRF are lower than those of conventional methods (Figure 8). For both methods, CV decreases as the acquisition duration increases. In case of *T*_1_ estimation by the MRF method, it takes 15 min to reach 10% of CV for all metabolites. On the other hands, it takes approximately 26 min to reach 10% of CV when using conventional methods. Considering *T*_2_ measurement, MRF shows 20% of CV within 15 min of acquisition time, which is comparable to multi-TE method.

**Figure 8:**
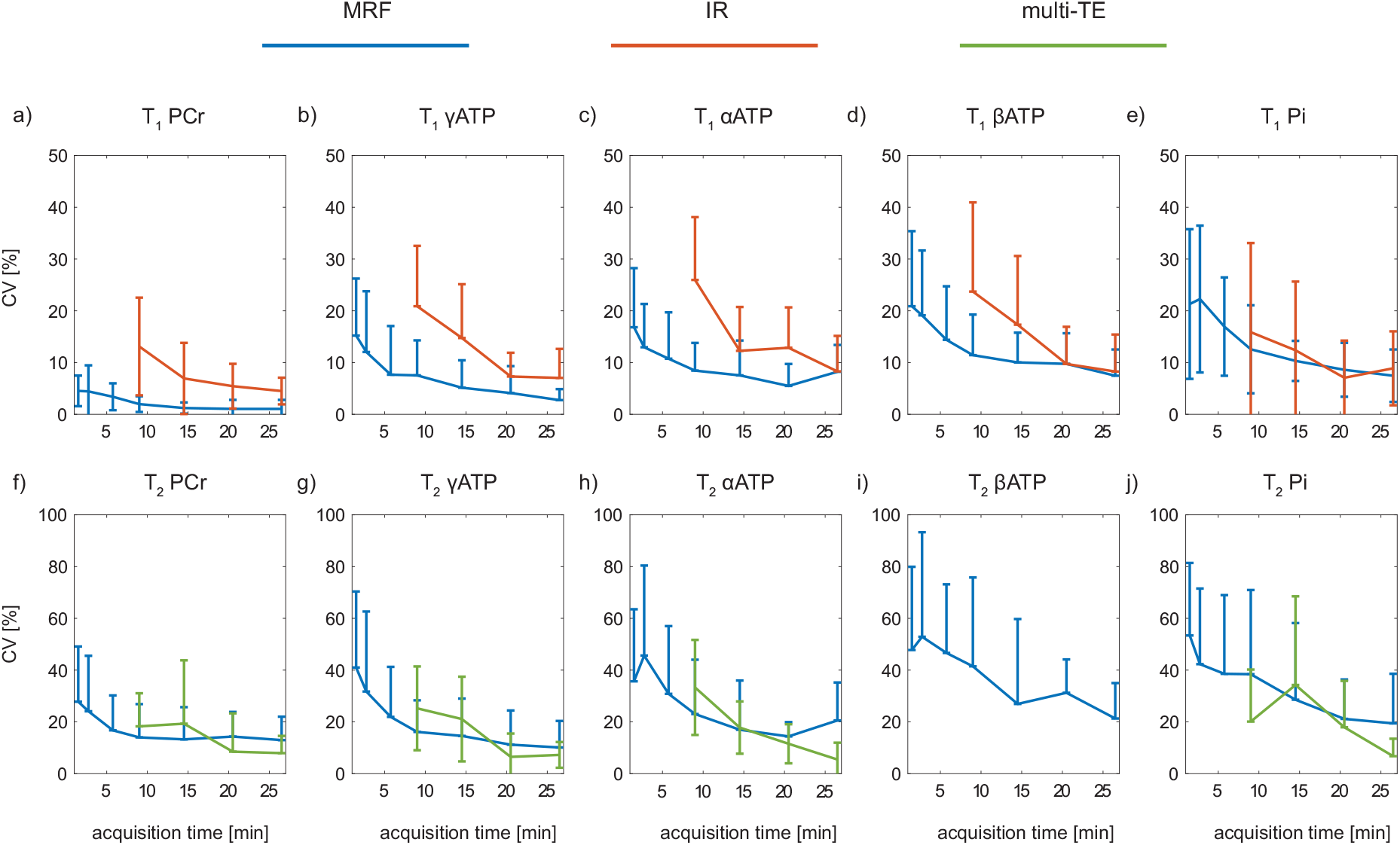
Evaluation of mean CV changes over acquisition time with SD. *T*_1_ and *T*_2_ estimated by MRF and conventional methods are shown in blue, orange, and green, respectively.

## Discussion

In the current study, fast measurements of *T*_1_ and *T*_2_ relaxation times of ^31^P metabolites by a bSSFP-based MRF framework has been introduced and demonstrated for the first time at 7T in human brain. This MRF scheme was evaluated using simulations and validated by *in vitro* experiments. The reproducibility and time efficiency of ^31^P MRF method were compared with the conventional inversion recovery (*T*_1_ measurement) and multi-TE (*T*_2_ measurement) methods *in vivo*, suggesting that MRF method can estimate *T*_1_ values of five metabolites with CV of less than 10% and *T*_2_ values of those with CV of less than 20% in 15 min with 40% of time reduction.

Measurements of *T*_1_ and *T*_2_ relaxation times of metabolites are usually time consuming using conventional methods due to the low signal intensity of ^31^P. MRF techniques are commonly based on bSSFP-type or gradient spoiled SSFP type sequences. The bSSFP sequence could attain high SNR and the obtained signal intensity is inherently sensitivity to *T*_1_ and *T*_2_ relaxation times. However, the bSSFP-type sequence introduces the commonly known banding artifact due to B_0_ field inhomogeneity (19). To avoid such periodical stopband and off-resonance sensitivity, a unbalanced stead-state type sequence was introduced for usual water signal based MRF studies (11). Considering the low sensitivity of ^31^P, the current MRF scheme employed the bSSFP-type sequence in order to take advantage of its merit in SNR.

Unlike MRF studies in MRI (8), where only one isochromat with a single off-resonance frequency could be assumed in small voxels, a distribution of off-resonance frequencies occurs in an 1D-slice. Therefore, an experimentally acquired *B*_0_ map was used to characterize the zero mean *B*_0_ distribution, which allows the accurate modeling of the bandpass induced in the bSSFP sequence and also reduces the complexity of the parameter estimation by eliminating one free parameter for dictionary matching. Stopband and passband (20) appear periodically, which is defined by the dephasing time within a given TR. In this initial approach, stopband has not been optimized for chemical shift ranges of all metabolites, as Pi peak is located near stopband, which results in suboptimal signal intensity of Pi as shown in supporting Figure (Figure S1). It is worth to mention that shorter TR can be preferable since less periods of the frequency response are needed to be taken into account to cover all metabolites at cost of a spectral resolution. Since the size of each voxel in 2D MRF is much smaller than a whole slice or non-localized spectroscopic MRF scheme, *B*_0_ map acquisition may not be needed but it can be estimated in the dictionary matching in the future.

Furthermore, there is significant *B*_1_ inhomogeneity caused by the surface coil (21). The effect of *B*_1_ inhomogeneity may not be substantial in small VOIs, while for a large VOI like a whole slice used in this study, this effect is not negligible. However, unlike acquiring *B*_0_ map, *B*_1_ mapping of ^31^P coil is time consuming and lengthens total acquisition duration because of the low sensitivity (22). Therefore, a *B*_1_ sensitive FA pattern (part B) (16) was added in our MRF sequence and *B*_1_ was then matched together with relaxation parameters.The phantom experiments also show that estimated *T*_1_ and *T*_2_ parameters by MRF were not affected by the input transmit voltage and matched *B*_1_ values corresponded to transmit voltages.

In our phantom validation results, matched *T*_1_ values by the MRF technique agree well with those acquired using the conventional method. The *in vivo* measurement also shows that *T*_1_ values of PCr, Pi, and ATPs measured by MRF are in good agreement with those obtained by the inversion recovery method. However, measured *T*_2_ values by MRF is shorter than those by multi-TE method. *T*_2_ underestimation has been reported in MRF studies using bSSFP readout (23; 24), which is mainly caused by intra-voxel phase dispersion especially at large *T*_2_. *T*_1_ underestimation is induced by slice profile imperfections (23; 24). Since slice profile was incorporated in the dictionary, measured *T*_1_ values between MRF and inversion recovery did not show statistical difference. In *in vivo* results estimated *T*_2_ values of PCr by MRF was much shorter than those obtained by multi-TE method, which could be partially resulting from the intra-voxle phase dispersion as described in previous research (23; 24). Besides the phase dispersion, magnetization transfer affects *T*_2_ estimation as well. Our model does not take into account the chemical exchange between PCr and *γ*ATP but considers all metabolites as an independent pool for both *in vitro* and *in vivo*. The supplementary figure shows the influence of chemical exchange inclusion on *T*_2_ value of PCr and *γATP* (Figure S3). There are some limitations in this preliminary study. Due to the passband filter effect, the Pi resonance falls into the stopband area, which affects its signal intensity. This can be improved in the future study by optimization of MRF acquisition parameters. Furthermore, our simulation results show that *T*_1_ estimation by MRF is more robust than *T*_2_ estimation, which is more sensitivity to 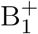. The incorporation of 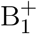 map or introducing a B_1_-insensitive acquisition scheme is expected to further improve the *T*_2_ estimation.

## Conclusions

In conclusion, our study shows the feasibility of fast relaxation time measurement in human brain by ^31^P spectroscopic MRF method. It enables to acquire *T*_1_ and *T*_2_ relaxation parameters within 15 min providing improved *T*_1_ measurement reproducibility compared to the conventional inversion recovery and multi-TE methods.

## Acknowledgments

This work was supported by the Swiss National Science Foundation (grants n^*°*^320030_189064). We acknowledge access to the facilities and expertise of the CIBM Center for Biomedical Imaging, a Swiss research center of excellence founded and supported by Lausanne University Hospital (CHUV), University of Lausanne (UNIL), Ecole polytechnique fédérale de Lausanne (EPFL), University of Geneva (UNIGE) and Geneva University Hospitals (HUG).

## Supporting Figures and Tables

**Table ST1:** Lower bounds of MRF dictionaries and their increment per step for simulation, *in vitro*, and *in vivo* data.

|  |  | lower bound |  | increment |  |
| --- | --- | --- | --- | --- | --- |
| | | $T_1$ [s] | $T_2$ [ms] | $T_1$ [s] | $T_2$ [%] |
| simu. | PCr | 3.10 | 30 | 0.05 | 1.05 |
|  | ATP | 0.72 | 20 | 0.02 | 1.04 |
| | | $T_1$ [s] | $T_2$ [ms] | $T_1$ [%] | $T_2$ [%] |
| *in vitro* | P1-P4 | 0.25 | 15 | 5.0 | 6 |
|  | P5-P7 | 2.10 | 30 | 4.2 | 6 |
| | | $T_1$ [s] | $T_2$ [ms] | $T_1$ | $T_2$ |
| *in vivo* | PCr | 3.0 | 30 | 3% | 6% |
| | γATP | 0.6 | 5 | 6% | 2 ms |
| | αATP | 0.65 | 5 | 6% | 2 ms |
| | βATP | 0.65 | 5 | 6% | 2 ms |
|  | Pi | 2.8 | 25 | 5% | 7% |

**Figure S1:**
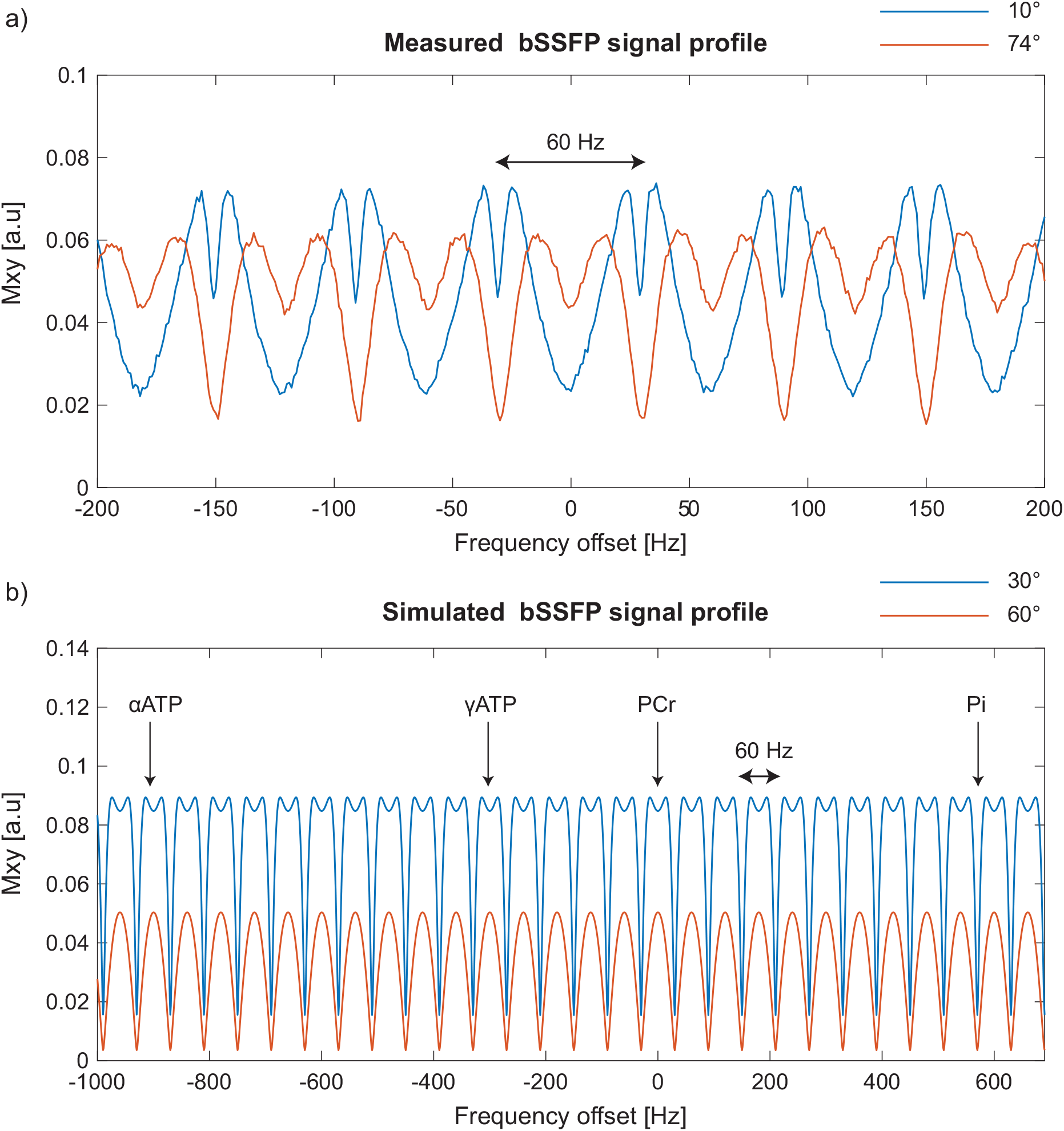
Measured and simulated bSSFP profiles. Measured bSSFP profiles with different flip angles are illustrated in (a). Simulated bSSFP profiles with different flip angles are presented in (b). Positions of PCr, *α*-, *γ*ATP, and Pi peaks are marked in (b).

**Figure S2:**
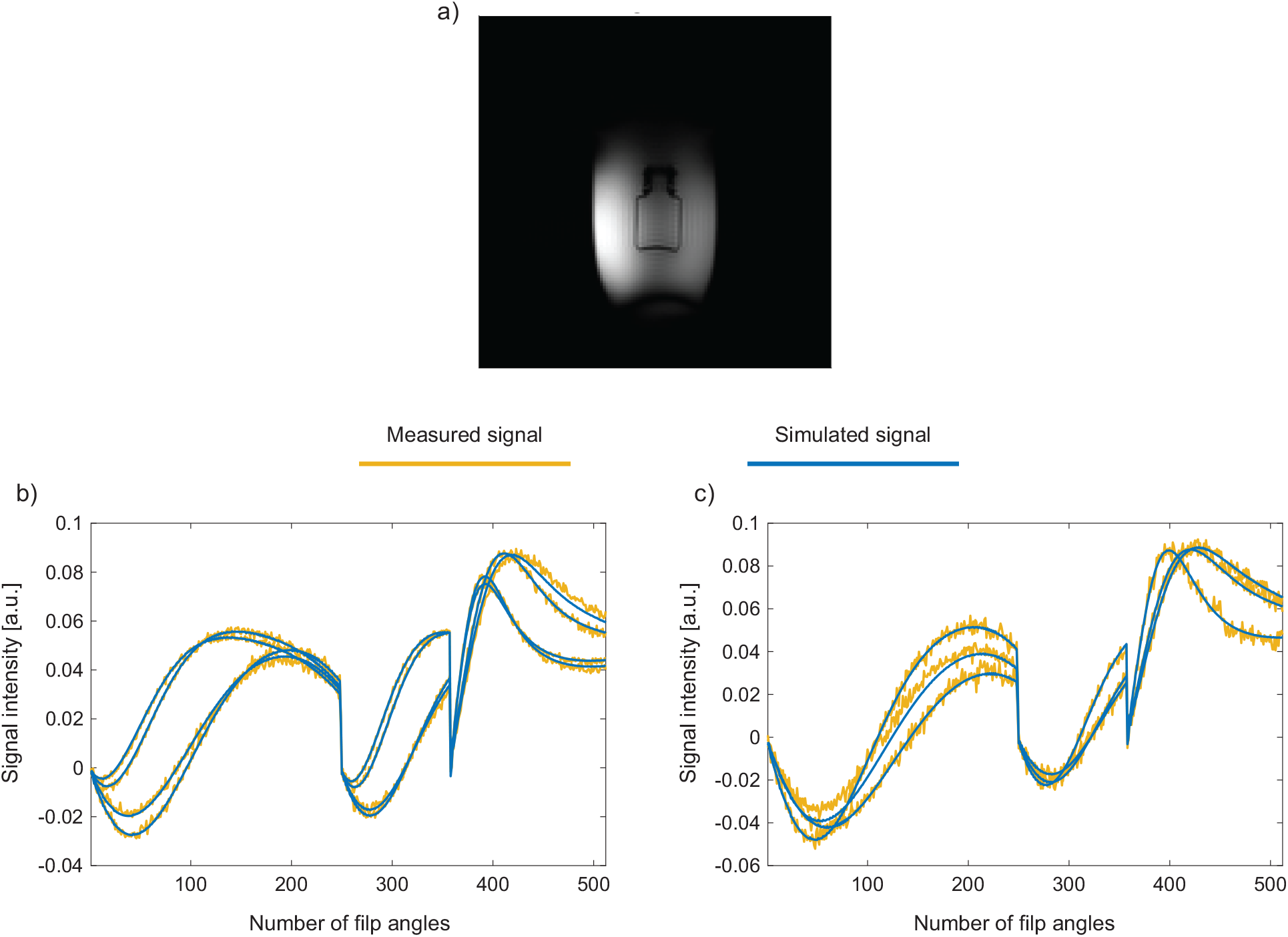
Representative reference image (a) and dictionary matching results (b-c). The outer bottle contains saline and inner bottle contains Pi and contrast agents. Measured signal from each phantom is illustrated in yellow and corresponding simulated signal in blue (b-c).

**Figure S3:**
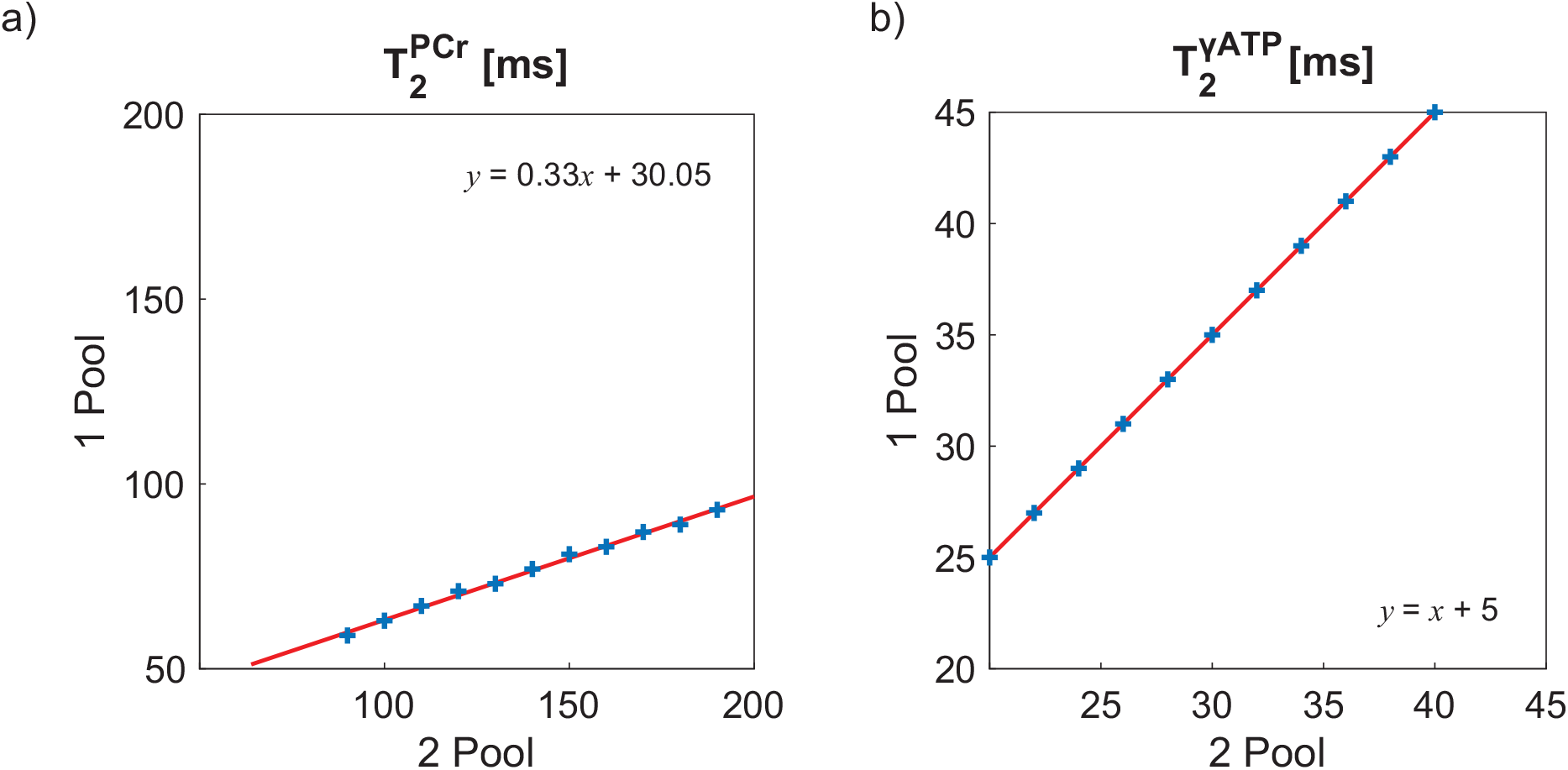
Estimated *T*_2_ relaxation times of PCr and *γ*ATP by 1 pool vs. 2 pool model. *T*_2_ values of PCr (a) and ATP (b) are presented in order. The y-axis indicates the matched *T*_2_ values by the single pool model and the x-axis shows the ground truth *T*_2_ values generated by 2 pool model. The linear regression lines are illustrated in red. The blue cross dots indicate estimated relaxation parameters by the two models.

The *T*_2_ matching error of the single pool model, the Bloch equations without a chemical exchange between PCr and *γ*ATP, was examined by fitting PCr and *γ*ATP signal evolutions generated using a 2 pool model, which is the Bloch-McConnell equations including creatine kinase rate. The 2 pool model simulation parameters were as follows. 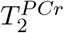 and 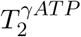 were altered in 10 ms steps between [90; 190] ms, keeping 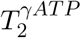 fixed to 30 ms, and in 2 ms steps in between [20; 40] ms, keeping 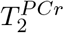 fixed to 135 ms, respectively. Other parameters were set as follows: 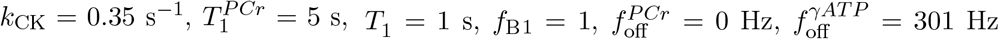, and 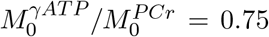. The generated signal evolutions were then fitted using dictionaries created by the single pool model. The matching results compared to the ground truth 2 pool values are shown in Figure S3. The matched 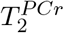 are linearly correlated with the 2 pool ground values, with an significant underestimation of around 50 % in the examined range. 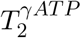 shows an offset leading to an overestimation of 5 ms.

## Notes

### Competing Interest Statement

The authors have declared no competing interest.

